# A novel SARS-CoV-2 related virus with complex recombination isolated from bats in Yunnan province, China

**DOI:** 10.1101/2021.03.17.435823

**Authors:** Li-li Li, Jing-lin Wang, Xiao-hua Ma, Jin-song Li, Xiao-fei Yang, Wei-feng Shi, Zhao-jun Duan

**Author notes:** **Corresponding author:** Dr. Zhao-jun Duan.

## Abstract

A novel beta-coronavirus, SARS-CoV-2, emerged in late 2019 and rapidly spread throughout the world, causing the COVID-19 pandemic. However, the origin and direct viral ancestors of SARS-CoV-2 remain elusive. Here, we discovered a new SARS-CoV-2-related virus in Yunnan province, in 2018, provisionally named PrC31, which shares 90.7% and 92.0% nucleotide identities with SARS-CoV-2 and the bat SARSr-CoV ZC45, respectively. Sequence alignment revealed that several genomic regions shared strong identity with SARS-CoV-2, phylogenetic analysis supported that PrC31 shares a common ancestor with SARS-CoV-2. The receptor binding domain of PrC31 showed only 64.2% amino acid identity with SARS-CoV-2. Recombination analysis revealed that PrC31 underwent multiple complex recombination events within the SARS-CoV and SARS-CoV-2 sub-lineages, indicating the evolution of PrC31 from yet-to-be-identified intermediate recombination strains. Combination with previous studies revealed that the beta-CoVs may possess more complicated recombination mechanism. The discovery of PrC31 supports that bats are the natural hosts of SARS-CoV-2.

## Introduction

Coronaviruses (CoVs) are a group of viruses that can infect humans and various mammalian and bird species (1, 2). So far, seven CoV species have been identified in humans. Of these, severe acute respiratory syndrome coronavirus (SARS-CoV) emerged in 2003 and caused multiple epidemics worldwide, and had a fatality rate of ~9.5%(3). Approximately ten years later, another highly pathogenic human CoV, Middle East respiratory syndrome coronavirus (MERS-CoV) emerged and caused numerous outbreaks in the Middle East and South Korea in 2015 (4–6). In December 2019, a novel beta-CoV, now termed severe acute respiratory syndrome coronavirus 2 (SARS-CoV-2), was first identified. SARS-CoV-2 caused a pneumonia outbreak in Wuhan, China, and eventually caused a pandemic, with > 116,521,000 reported cases and > 2,589,000 deaths worldwide as of March 9, 2021 (7–10).

Both SARS-CoV and MERS-CoV are likely to have originated from bats (5, 10–13). Many SARS-related coronaviruses (SARSr-CoV) have been discovered in bats following CoV outbreaks (11, 14–16), suggesting that bats may be the natural hosts of SARS-CoV. Similarly, several MERS-related coronaviruses have also been islolated from various bat species (5). Notably, palm civets and dromedary camels most likely served as intermediate hosts for SARS-CoV and MERS-CoV, respectively, because these animals carried almost identical viruses to the SARS-CoV and MERS-CoV strains isolated from humans (5). Furthermore, two human coronaviruses, HCoV-NL63 and HCoV-229E, are also considered to have originated in bats, whereas HCoV-OC43 and HKU1 were likely to have originated from rodents (5, 17).

Since the identification of SARS-CoV-2, CoVs phylogenetically related to SARS-CoV-2 (RaTG13, RmYN02, Rc-o319, RshSTT182, RshSTT182200 and RacCS203) have been discovered in bats from China, Japan, and Cambodia (7, 18–23), with most of them discovered by analyzing stored frozen samples (7, 10, 14, 19–22). Of these, RaTG13 and RmYN02, which were identified in Yunnan province, China, shared whole-genome nucleotide sequence identities of 96.2% and 93.3% with SARS-CoV-2, respectively (7, 19). SARS-CoV-2-related CoVs were also identified in pangolins, whose receptor binding domain (RBD) shared up to 97.4% nucleotide identity with that of SARS-CoV-2 (20, 21). This suggests that pangolins are a potential host of SARS-CoV-2, although the role of pangolins in the evolutionary history of SARS-CoV-2 remains elusive. Nevertheless, either the direct progenitor of SARS-CoV-2 is yet to be discovered, or the transmission route of SARS-CoV from bats to humans via an intermediate host must still be determined (24). The discovery of more SARS-CoV-2-related viruses will help to clarify the details regarding the emergence and evolutionary history of SARS-CoV-2.

## Results

### Identification of a novel SARS-CoV-2-related coronavirus

Based on the molecular identification results, all collected bats belonged to five different species: *Rhinolophus affinis, Miniopterus schreibersii, Rhinolophus blythi, Rhinolophus pusillus*, and *Hipposideros armiger*. By retrospectively analyzing our NGS data, we found a new bat beta-CoV related to SARS-CoV-2 in *Rhinolophus blythi* collected from Yunnan province, China, in 2018. The qRT-PCR results revealed that two samples tested positive for SARS-CoV-2 with *Ct* values of 32.4 (sample C25) and 35.6 (sample C31). Both bats were identified as *Rhinolophus blythi*. A near complete genome of this virus comprising 29,749 bp was obtained from sample C31 and tentatively named PrC31. The virus genome isolated from the second positive sample had the same sequence as PrC31.

### Genetic characteristics and comparison with SARS-CoV-2 and other related viruses

Analysis of the complete PrC31 genome revealed that it shared 90.7% and 92.0% nucleotide identity to SARS-CoV-2 and bat SARSr-CoV ZC45, respectively (Table 1). Although the whole genome of PrC31 was more closely related to ZC45 compared to the other viruses examined, several genes of PrC31 showed highly similar nucleotide identities (> 96%) with SARS-CoV-2, including E, ORF7a, ORF7b, ORF8, N and ORF10 (Table 1). Notably, ORF8 and ORF1a (the region spanning nucleotides 1-12719) of PrC31 were genetically closer to SARS-CoV-2 than any other viruses identified to date, exhibiting 98.1% and 96.6% nucleotide identities, respectively. However, in other regions, PrC31 was more similar to SARS-CoV or SARSr-CoV ZC45.

**Table 1:** Sequence identities comparing PrC31 with SARS-CoV-2 and other representative beta-CoVs

|  |  | Complete | Genes |  |  |  |  |  |  |  |  |  |  |  |
| --- | --- | --- | --- | --- | --- | --- | --- | --- | --- | --- | --- | --- | --- | --- |
| Strain |  | Genome | 1ab | S | RBD | 3a | E | M | 6 | 7a | 7b | 8 | N | 10 |
| nt | Wuhan-Hu-1 | 90.7 | 92.6 | 74.9 | 61.5 | 89.2 | 99.1 | 93.4 | 94.4 | 96 | 97.8 | 98.1 | 97 | 99.1 |
| (%) | RaTG13 | 90.4 | 92 | 75.6 | 61.6 | 88.7 | 98.7 | 93.3 | 95.9 | 93.8 | 98.5 | 97.8 | 96.6 | 98.3 |
|  | ZC45 | 92 | 91.2 | 94.8 | 95.3 | 95.6 | 98.7 | 96 | 91.8 | 87.9 | 95.6 | 89.4 | 91.3 | 98.3 |
|  | RmYN02 | 90.4 | 93.2 | 74.3 | 81.1 | 88.5 | 97.8 | 91 | 95.4 | 95.4 | 93.3 | 48.8 | 98.1 | 98.3 |
|  | Pangolin/GXP5L | 83.3 | 83.5 | 75.2 | 61.9 | 85.1 | 96.5 | 90.9 | 89.3 | 85.5 | 84.4 | 81 | 91.2 | 94.9 |
|  | Pangolin/GD | 87.9 | 90.3 | 79.2 | 61.2 | 90.6 | 98.3 | 93.1 | 91.3 | 91.7 | 94.1 | 93.2 | 96.2 | 98.3 |
|  | Rc-o319 | 79.3 | 80.3 | 70.7 | 63.8 | 79.3 | 96.5 | 84.8 | 85.2 | 77.2 | 79.3 | 46.3 | 87.7 | 95.7 |
|  | Rs4237 | 82.3 | 83.2 | 74.5 | 82.1 | 75.6 | 92.2 | 82.7 | 76.1 | 82.8 | 80.7 | 65.8 | 87.9 | 92.3 |
|  | Tor2 | 81.4 | 83 | 71.6 | 63.7 | 74.5 | 92.6 | 83.4 | 74.6 | 80.6 | 80.7 | 44.6 | 88.1 | 93.2 |
|  | RShSTT182 | 88.6 | 90.3 | 72.2 | 62.9 | 87.6 | 98.3 | 90.0 | 87.6 | 94.4 | 99.3 | 95.1 | 94.3 | 98.3 |
|  | RacCS203 | 88.8 | 90.3 | 74.4 | 80.1 | 87.8 | 98.2 | 92.5 | 91.4 | 90.8 | 93.2 | 92.9 | 93.7 | 99.1 |

The RBD of PrC31 was evolutionarily distant from SARS-CoV-2, sharing only 64.2% amino acid identity, whereas it was almost identical to that of ZC45, with only one amino acid difference. Similar to most bat SARSr-CoVs, one long (14 aa) deletion and one short (5 aa) deletion were present in PrC31, which were absent from SARS-CoV, SARS-CoV-2, pangonlin-CoV and RaTG13. We predicted the three-dimensional structure of the RBD of PrC31, ZC45 and SARS-CoV-2 using homology modeling. Similar to RmYN02, the two loops close to the receptor binding site of the PrC31 RBD were shorter than those of SARS-CoV-2, due to two deletions; this region may influence the binding capacity of the PrC31 RBD with the angiotensin converting enzyme 2 (ACE2) receptor (Fig.1A–1D). Moreover, of the six amino acid residues that are essential for the binding of the SARS-CoV-2 spike protein to ACE2 (L455, F486, Q493, S494, N501, and Y505), PrC31 and RmYN02 possesed only one(Y505) (Fig.1E)

**Figure 1:**
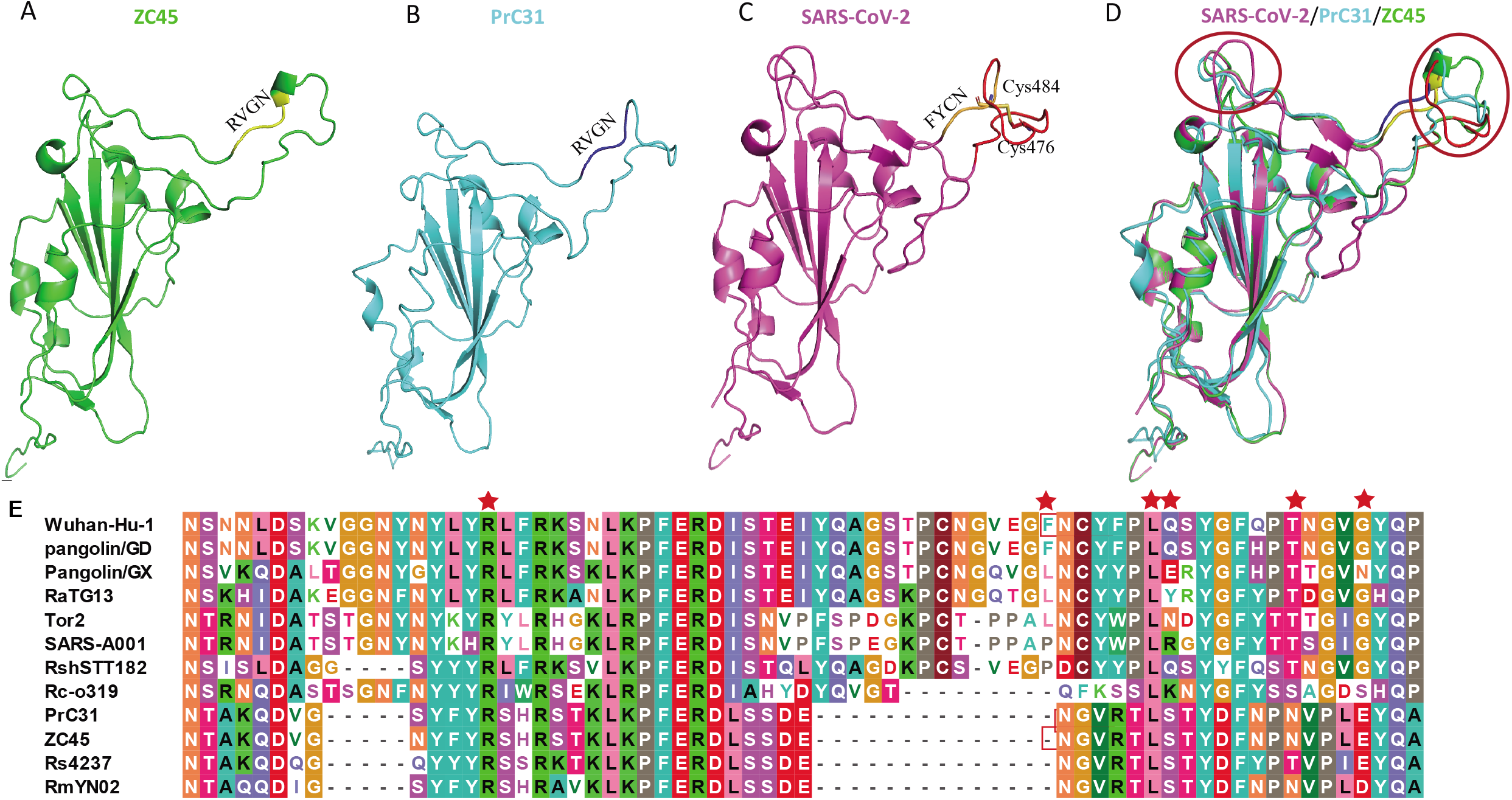
Homology modeling structures and Characterization of Receptor binding domain (RBD) of PrC31 and Representative Beta-CoVs. (A-D) Homology modeling structures of PrC31 and Representative Beta-CoVs. The three-dimensional structures of PrC31, ZC45 and SARS-CoV-2 RBDs were modeled using the Swiss-Model program, using the SARS-CoV-2 RBD structure (PDB: 7a91.1) as a template. The two deletion loops in PrC31 and ZC45 are marked with a circle. (E) Characterization of the RBDs of PrC31 and representative beta-CoVs. The six critial amino acid residues for ACE2 interaction were marked using red star.

### Phylogenetic analysis of PrC31 and representative sarbecoviruses

Phylogenetic analysis of the complete PrC31 genome revealed that it belonged to a separate clade to SARS-CoV-2, while most other SARS-CoV-2-related viruses were grouped together (Fig.2). However, the PrC31 RNA-dependent RNA polymerase was phylogenetically grouped within the SARS-CoV lineage and clustered with bat SARS-rCoV. The spike protein of PrC31 fell within the SARS-CoV-2 sub-lineage and clustered with ZC45 and CXZ21, while being distant from SARS-CoV-2. The topological differences between various regions of PrC31 strongly suggest the occurrence of recombination events throughout its evolution.

**Fig. 2.**
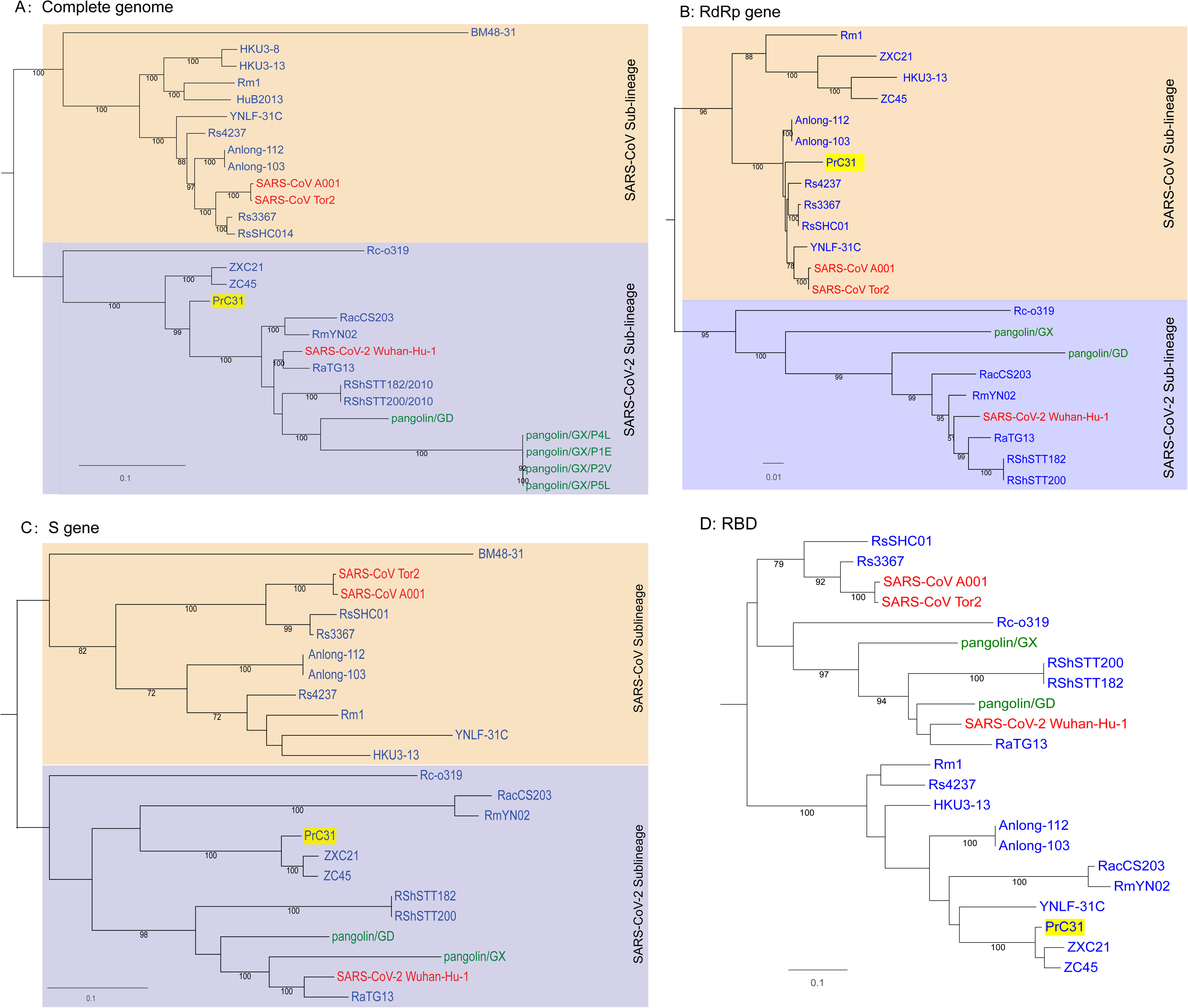
Phylogenetic trees of SARS-CoV-2 and representative sarbecoviruses. (A)Complete genome; (B)RdRp gene (C)S gene (D)RBD region. SARS-CoV lineage and SARS-CoV-2-related lineages are shown in orange and purple shadow, respectively. Viruses that originated in bats are labeled in blue, human viruses are labeled in red and pangolin viruses are labeled in green. The PrC31 identified in this study is highlighted in yellow shadow. Phylogenetic analyses were performed with RaxML software (v8.2.11) using the GTR nucleotide substitution model, GAMMA distribution of rates among sites, and 1000 bootstrap replicates

### Multiple and complex recombination events in the evolution of PrC31

The full-length genome sequences of PrC31 and closely related beta-CoVs were aligned to search for possible recombination events. Strikingly, both the similarity and bootstrap plots revealed multiple and complex long-segment recombination events in PrC31, which likely arose from multiple beta-CoVs from within the SARS-CoV and SARS-CoV-2 sub-lineage. As shown in Figure 3, three recombination breakpoints were detected. For the region spanning nucleotides 1-12,719 and 27,143 to the 3’ terminus of the genome, PrC31 was most closely related to SARS-CoV-2 and RmYN02. In these regions, PrC31 was phylogenetically grouped with RmYN02 and in a sister clade to SARS-CoV-2 (Figure 4a and 4d). For the 12,720-20,244 nucleotide region, which included ORF1ab, PrC31 was grouped with SARS-CoV and bat SARSr-CoVs (Figure 4b). Moreover, PrC31 presented the highest similarity to ZC45 in the 20,245-27,142 genomic fragment, which included part of ORF1ab, S, ORF3, E, and part of the M gene, and fell within the SARS-CoV-2 sub-lineage (Figure 4c)

**Fig. 3.**
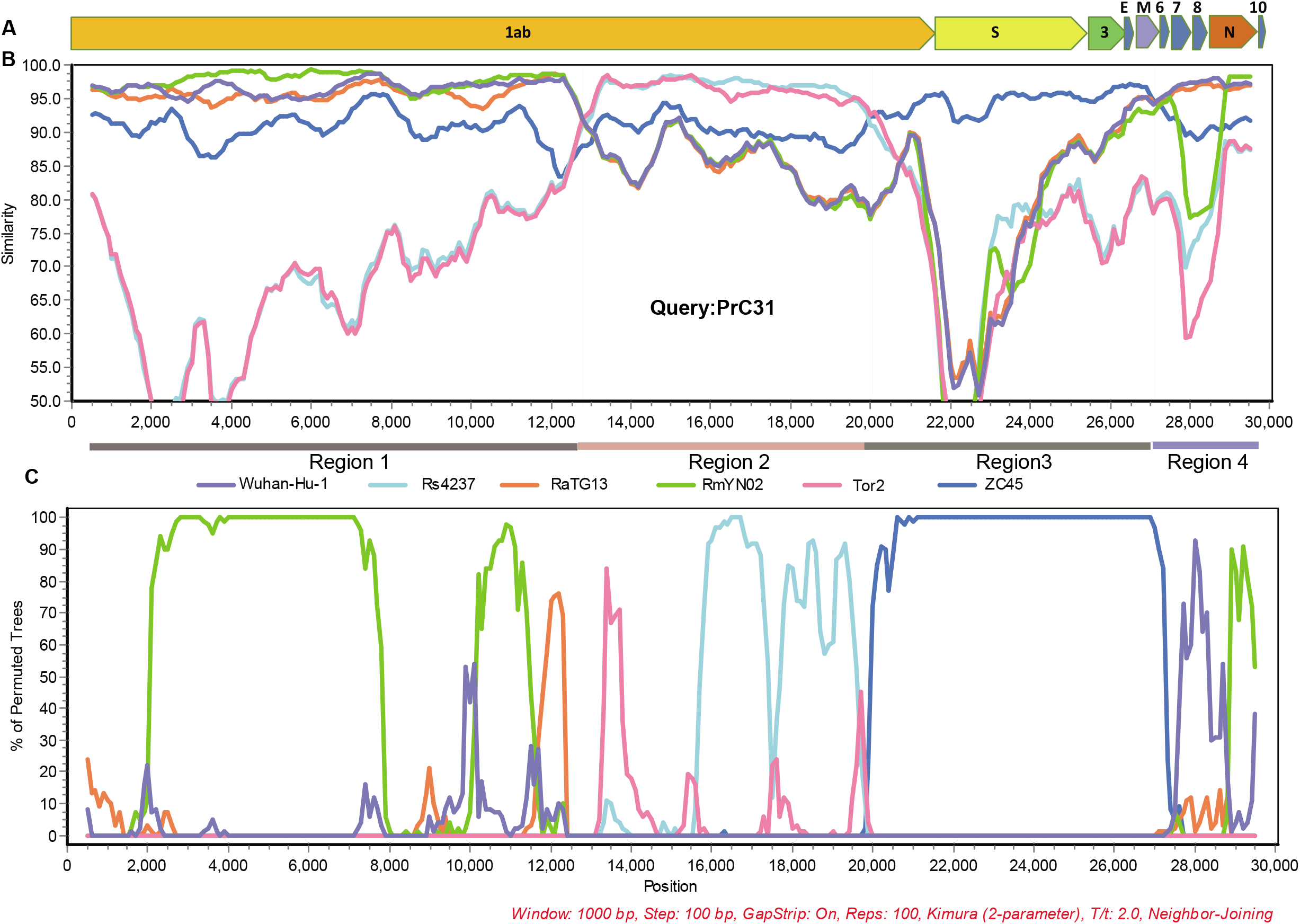
Recombination analysis. A. Genome organization of PrC31. (B) Similarity plot and (C) Bootstrap plot of full-length genome of human SARS-CoV-2, pangolin- and bat beta-CoVs using PrC31as the query. Slide window was set to 1000 bp with 100 bp steps.

**Fig. 4.**
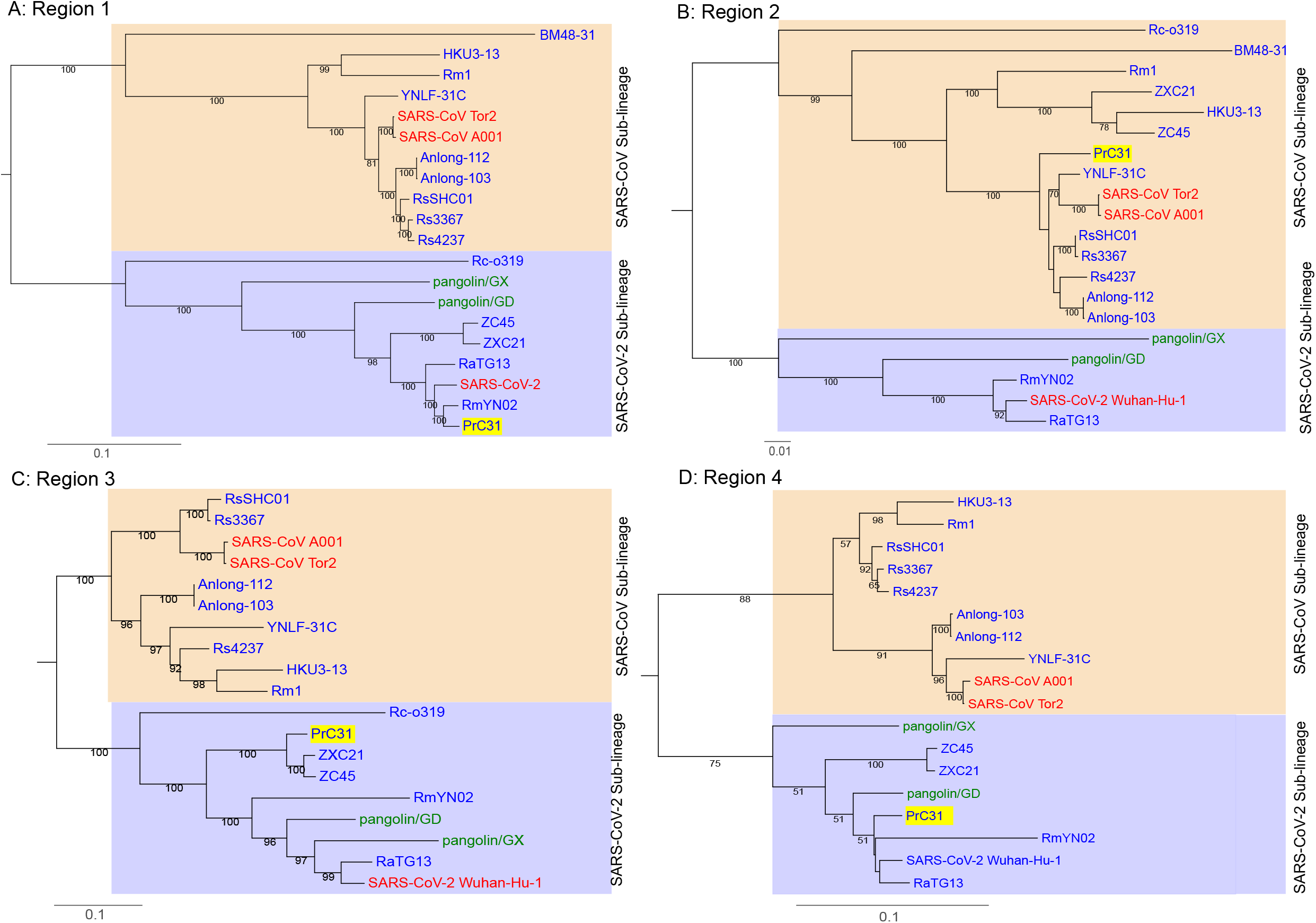
Phylogenetic trees of various regions of the PrC31 genome. SARS-CoV and SARS-CoV-2-related lineages are shown as orange and purple shadow, respectively. The PrC31 virus identified in this study is indicated with yellow shadow. Viral taxonomy is labeled in color that originated in bats are labeled in blue, humans in red, and pangolins in green. Phylogenetic analyses were performed with RaxML software (v8.2.11) using the GTR nucleotide substitution model, GAMMA distribution of rates among sites, and 1000 bootstrap replicates

## Discussion

The recently-emerged SARS-CoV-2 virus triggered the ongoing COVID-19 pandemic, which has high morbidity and fatality rates, and poses a great threat to global public health. Identifying the origin and host range of SARS-CoV-2 will aid in its prevention and control, and will facilitate preparation for future CoV pandemics. Although several SARS-CoV-2-related viruses were detected in bats and pangolins, none of them appear to be the immediate ancestor of SARS-CoV-2; the exact origin of SARS-CoV-2 is still unclear (12, 25). In this study, we discovered PrC31, a sarbecovirus isolated from bat intestinal tissues collected in 2018. PrC31 phylogenetically falls into the SARS-CoV-2 clade and has undergone multiple and complex recombination events.

Animals that continuously harbor viruses closely related to SARS-CoV-2 for extended time periods can become natural SARS-CoV-2 hosts (2). To date, several bat viruses have been identified that have strong sequence similarities to SARS-CoV-2, sharing more than 90% sequence identity. Especially, RaTG13 possesses 96.2% identity with SARS-CoV-2 (7, 18, 19, 21, 23). The PrC31 virus identified in this study showed 90.7% genome identity with SARS-CoV-2; notably, the E, ORF7, ORF8, N and ORF10 genes shared more than 96% identity with SARS-CoV-2. Both the genetic similarity and diversity of SARS-CoV-2-related viruses support the claim that bats were the natural hosts of SARS-CoV-2 (10, 19).

Recombination events between various SARSr-CoVs have occurred frequently in bats (5, 16). SARS-CoV-2 may also be a recombined virus, potentially with the backbone of RaTG13 and a RBD region acquired from pangolin-like SARSr-CoVs (12, 21). In this study, we found that PrC31 phylogenetic clustered with SARS-CoV-2 and its related viruses. The results from our phylogenetic analyses suggested that recombination had occurred in PrC31. The similarity plot indicated that the PrC31 was subjected to multiple and complex recombination events involving more than two sarbecoviruses in the SARS-CoV and SARS-CoV-2 sub-lineages. The three breakpoints of PrC31 separate the genome into four regions. Region 1 (within ORF1a) and region 4 of PrC31 were closely related to SARS-CoV-2, RaTG13 and RmYN02. Region 2 of PrC31 was more similar to members of the SARS-CoV sub-lineage, including SARS-CoV and SARSr-CoV Rs4237 strain; region 3 was more closely related to ZC45 within SARS-CoV-2 sub-lineage. The multiple recombination events of PrC31 hint toward the existence of intermediate recombination strains within the SARS-CoV and SARS-CoV-2 sub-lineages that are yet to be identified. Our work suggests that the backbone of PrC31 may have evolved from a recent common ancestor of RaTG13, RmYN02 and SARS-CoV-2, and that it acquired regions 2 and 3 from precursor viruses of SARS-CoV and SARSr-CoV ZC45, respectively.

At present, the precise patterns and mechanisms driving recombination in sarbecoviruses are largely unknown. A recent report identified 16 recombination breakpoints in 69 sarbecoviruses (26), although in the majority of strains, the recombination sites were located within the S gene and upstream of ORF8 (5, 9, 16). The three recombination breakpoints of PrC31 were located in ORF1a, ORF1b and M genes with long fragment recombination, suggestive of a complicated recombination pattern in sarbecoviruses. Similar to PrC31, SARS-CoV-2 may have evolved via complex recombination between various related coronaviruses or their progenitors (10). In fact, the direct progenitor of SARS-CoV may have evolved by recombination with progenitors of SARSr-CoV (Hu et al. 2017). Together, these findings suggest that recombination and its role in the evolution history of sarbecoviruses may be more complicated and significant than initially expected.

Pangolins may also harbor ancestral beta-CoVs related to SARS-CoV-2 (2, 20, 21); the receptor-binding motif of pangolin beta-CoVs share an almost identical amino acid sequence with SARS-CoV-2 (20, 21), suggesting that SARS-CoV-2 may have acquired its RBD region from a pangolin CoV via recombination(27). However, unlike bats, pangolins infected with beta-CoVs present overt symptoms and eventually die, rendering them unlikely to be natural hosts. Intermediate hosts generally serve as zoonotic sources for human infection, acting as vectors for viral replication and transmission to humans (2). Current evidence suggests that pangolins were not the direct intermediate hosts of SARS-CoV-2. However, pangolins certainly played an important role in the evolutionary history of SARS-CoV-2 related viruses, eventually leading to the transmission of SARS-CoV-2 to humans. It cannot be excluded that a novel recombination event involving SARS-CoV-2 or SARS-CoV-2 related viruses and SARS-CoV or SARSr-CoV will lead to the virus presumed as “SARS-CoV-3”, which may be transmitted to human populations in the future.

The discovery of PrC31 provides more evidence for the bat origin of SARS-CoV-2 (10, 28). Identifying more SARS-CoV-2 related viruses in nature will provide deeper insight into the origins of SARS-CoV-2. It will be necessary to expand the sampling areas and animal species examined to find more close relatives of SARS-CoV-2. There may be an unknown intermediate host of SARS-CoV-2 that played a similar role to that of civets and camels in the SARS-CoV and MERS-CoV epidemics, respectively. Furthermore, PrC31 was firstly tested for positive using SARS-CoV-2 qPCR kit, which targets the ORF1ab and N genes of SARS-CoV-2. This emphasizes the need to gather sequence information for positive samples during environmental surveillance of SARS-CoV-2, as samples may be contaminated with a closely related beta-CoVs from wild animals such as bats.

## Materials and methods

We retrospectively analyzed bat next generation sequencing (NGS) data that we performed in 2019, and found SARS-CoV-2-related reads present in one pool of intestinal tissues. The details of sampling and high-throughput sequencing are given below.

### Sample collection and pretreatment

In 2018, 36 bats were captured in Yunnan province, China. The bats were dissected following anesthetization. Liver, lung, spleen and intestinal tissue specimens were collected and transported to the Chinese center for disease control, where they were stored at −80°C until further analysis. The bat species were identified by polymerase chain reaction (PCR) to amplify the cytochrome B gene, as previously described (29). Intestinal tissues collected from 36 bats were homogenized in minimum essential medium and the suspensions were centrifuged at 8,000 rpm. The supernatants were merged into two pools according to bat species, then digested using DNase I for RNA Extraction. All procedures were performed in a biosafety cabinet in a biosafety level 2 facility. This study was approved by the ethics committee of the CCDC, and was performed according to Chinese ethics, laws and regulations.

### RNA extraction and next-generation sequencing (NGS)

Nucleic acids were extracted using a QIAamp MinElute Virus Spin Kit (QIAGEN) and used to construct the sequencing libraries. The library preparation and sequencing steps were performed by Novogene Bioinformatics Technology (Beijing, China). In brief, the ribosomal RNA was removed using the Ribo-Zero-Gold (Human–Mouse–Rat) Kit (Illumina, USA) and the Ribo-Zero-Gold (Epidemiology) Kit (Illumina). The libraries were constructed using a Nextera XT kit (Illumina), and sequencing was performed on the Illumina NovaSeq 6000 platform according to the procedure for transcriptome sequencing.

### Bioinformatic analyses

Bioinformatics analysis of the sequencing data was conducted using an in-lab bioinformatics analysis platform. Prinseq-lite software (version 0.20.4) was used to remove lower quality reads, and Bowtie2 was used to align and map the filtered reads to the host reference genome. Mira (version 4.0.2) was used for *de novo* assembly of the clean reads. Both BLASTn and BLASTx of the BLAST+ package (version 2.2.30) were used to search against local viral nucleotide and protein databases. The E-value cut-off was set to 1 × 10^−5^ to maintain high sensitivity and a low false-positive rate when performing BLAST searches.

### Sequencing of full-length genomes and quantitative real-time PCR (qRT-PCR)

We obtained reads that showed 96-98% nucleotide identity to SARS-CoV-2 from the PrC31 genome library. To confirm the sequences obtained from NGS and to fill the gaps, we designed 32 primer pairs according to the consensus sequences from the NGS and the conserved regions of SARS-CoV-2, RaTG13 and RmYN02, to amplify the whole PrC31 genome with at least 100 bp overlap between adjacent PCR fragments (Table S1). The PCR products were subjected to Sanger sequencing with pair-end sequencing. The 25 bp at the 5’ and 3’ termini were omitted, and the remaining sequences were assembled using Geneious Prime. Positive samples were quantified using TaqMan-based qPCR kit targeting the ORF1ab and N genes (BioGerm, China).

### Phylogenetic and recombination analyses

The complete genome sequences of reference viruses were downloaded from GenBank (https://www.ncbi.nlm.nih.gov/) and GISAID (https://www.gisaid.org/). The complete genome of PrC31 was aligned with representative SARS-CoV, SARS-CoV-2 and SARSr-CoV using Mafft (v7.475). Phylogenetic analyses were performed with RaxML software (v8.2.11) using the general time reversible nucleotide substitution model, GAMMA distribution of rates among sites, and 1000 bootstrap replicates. Potential recombination events were screened using RDP4 software and further analyzed by similarity plot using Sìmplot (v3.5.1) with potential major and minor parents.

### Structural modeling

The three-dimensional structures of PrC31, ZC45 and SARS-CoV-2 RBDs were modeled with the Swiss-Model program using the SARS-CoV-2 RBD structure (PDB: 7a91.1) as the template.

## Data availability

The sequences of PrC31 generated in this study were deposited in the GISAID and GenBank databases with the accession numbers EPI_ISL_1098866 and MW703458, respectively.

## Acknowledgements

This study was supported by National Science and Technology Major Project of China (No.2018ZX10305409-004-002)

## Author contributions

L-L L, Acquisition of data, Analysis and interpretation of data, Conception and design, Drafting or revising the article; M-XH, Acquisition of data; J-S L, Conception and design experiment; J-L W, Sample collection, Acquisition of data. W-F S, Analysis and interpretation of data, Conception and design, Drafting or revising the article. Z-J D, Conception and design, Analysis and interpretation of data, Drafting or revising the article.

