## Supplemental Table 1 for "A novel SARS-CoV-2 related virus with complex recombination isolated from bats in Yunnan province, China"

Supplemental Table 1 The primers used to in this study

| Primers (nucleotide positions) | Sequence, 5’→3’ | Product length, bp |
| --- | --- | --- |
| CV-F1（1-20） | ACAAACCAACTAACTCTCG | 306 |
| CV-R1（288-306） | TCTGATAAAGCCTCCTCCA |  |
| CV-F2（105-122） | CAGGTTGCTTACGGTTTC | 703 |
| CV-R2（790-807） | AGATCTTCTAGCTCGTGC |  |
| CV-F3（493-512） | ACTTGGTGTCCTTGTCCCTC | 927 |
| CV-R3（1400-1419） | GAGAACACACAGCCTCCAAA |  |
| CV-F4（1255-1274） | TGGTTACCTACCCCAAAATG | 1446 |
| CV-R4 （2681-2700） | ACACTCTTGTAACCTTGCAC |  |
| CV-F5（2242-2261） | GGCTTTGTGTGCTGACTCTA | 1463 |
| CV-R5（3686-3704） | GTTCGCACAAATGTCTACC |  |
| CV-F6（3307-3326） | TGCAGACATTGTGGAAGAAG | 738 |
| CV-R6（4025-4044） | CCCACTATATATGGAGCATC |  |
| CV-F7（3686-3705） | GTTCGCACAAATGTCTACCT | 894 |
| CV-R7（4560-4579） | CATGTACCGAGCAGCTTCTT |  |
| CV-F8（4283-4300） | CTTGGAACTGTTTCTTGG | 1206 |
| CV-R8（5471-5488） | CATAACAGCTTCTACACC |  |
| CV-F9（5056-5073） | CACTTTACGTGCTGAGGC | 1171 |
| CV-R9（6207-6226） | GCTCCAAAGGCAACGTATAC |  |
| CV-F10（5595-5614） | TTATGATGTCAGCACCACCT | 1969 |
| CV-R10（7545-7563） | CCAGCACAGAATGTATCAC |  |
| CV-F11（7040-7059） | CCTTGTAGTGTTTGTCTTAG | 1078 |
| CV-R11（8098-8117） | CATCTGAATCAACAAACCCT |  |
| CV-F12（7917-7936） | ATGTTGGTGATAGTGCGGAA | 2388 |
| CV-R12（10285-10304） | AAGTCTGTCCTGGTTGAATG |  |
| CV-F13（10182-10200） | GTAATGTTCAACTCAGGGT | 993 |
| CV-R13（11156-11174） | TCATAATACGCATCACCCA |  |
| CV-F14（10717-10731） | ACTAGGACCTCTTTCTGCTC | 969 |
| CV-R14（11666-11685） | ATGCTATTCTTTGGTGGGAG |  |
| CV-F15（11142-11160） | ATATGCCTGCTAGTTGGGT | 1053 |
| CV-R15（12176-12195） | GCCATCTTTTCCAACTTACG |  |
| CV-F16（11713-11732） | GTTGGGTGTTGGAGGTAAAC | 673 |
| CV-R16（12366-12385） | GGCTGCTGTTGTAAGGGGTA |  |
| CV-F17（12064-12081） | AGAAGCTTATGAGCAGGC | 1268 |
| CV-R17（13313-13331） | CACAACTACAGCCATAACC |  |
| CV-F18（13130-13149） | GATCAAGAATCCTTTGGTGG | 1465 |
| CV-R18（14576-14595） | ACCGGGTTTGACAGTTTGAA |  |
| CV-F19（13175-13192） | TGCCACATAGATCATCCA | 1928 |
| CV-R19（15084-15102） | AGTGGCGGCTATTGATTTC |  |
| CV-F20（14924-14941） | ATCAAGATGCACTTTTCG | 1652 |
| CV-R20（16558-16575） | AGTAGCTTTGAGCGTTTC |  |
| CV-F21（16462-16481） | ACTGACTTCAATGCGATAGC | 1310 |
| CV-R21（17752-17771） | TCTGAACCCTGTGATGAATC |  |
| CV-F22（17374-17392） | CCACGCACATTGCTAACTA | 2553 |
| CV-R23（19908-19926） | TGTCCTTCCACTCTACCAT |  |
| CV-F24（19793-19813） | CTGGGACTACAAAAGAGATGC | 1058 |
| CV-R24（20830-20850） | CGTAGGCAACCACTGTCTTA |  |
| CV-F25（20571-20590） | TACAATCTAGTCAAGCGTGG | 1777 |
| CV-R25（22330-22347） | GTCAAGTGCACAGTCTAC |  |
| CV-F26（22202-22220） | CAATTCACAGAGGAGACCC | 2025 |
| CV-R26（24207-24226） | CACTGGCTGTGGATGTCAAA |  |
| CV-F27（23981-23998） | CCACCTTTGCTCACAGAT | 1207 |
| CV-R27（25166-25187） | CTGGCTCAGAGTCGTCTTCATC |  |
| CV-F28（25038-25058） | GGTATGTTTGGCTTGGCTTCA | 1492 |
| CV-R28（26510-26529） | AAAGCAGCCAGAGGAAGATT |  |
| CV-F29（26034-26053） | GACTACTAGCGTGCCTTTGT | 1622 |
| CV-R29（27636-27655） | ACAAGGAATAGCAGAAAGGC |  |
| CV-F30（26952-26969） | TACAGGATTGGCAACTAC | 1084 |
| CV-R30（28016-28035） | AACGCACTACAAGACTACCC |  |
| CV-F31（27635-27654） | AGCCTTTCTGCTATTCCTTG | 969 |
| CV-R31（28584-28603） | TCCTTGAGGAAGTTGTAGCA |  |
| CV-F32（27830-27848） | TGATGACCCGTGTCCTATT | 1812 |
| CV-R32（29623-29641） | TAGGGCTCTTCCATATAGG |  |
